# Fatty acid composition and parasitism of European sardine (*Sardina pilchardus*) and European anchovy (*Engraulis encrasicolus*) populations in the northern Catalan Sea in the context of changing environmental conditions

**DOI:** 10.1101/2020.06.15.153775

**Authors:** Sebastian Biton-Porsmoguer, Ricard Bou, Elsa Lloret, Manuel Alcaide, Josep Lloret

## Abstract

The status of sardine and anchovy populations in the northern Mediterranean Sea has been declining in recent decades. In this study, fatty acids and parasitism at different reproductive and feeding stages in these two species were assessed using specimens caught along the northern Catalan coast, in order to assess the links between lipid dynamics, reproduction and feeding in these two species, and to contribute towards an explanation of the potential causes of the current situation of the stocks. The results support the use of fatty acid levels as indicators of the body condition of sardine and anchovy at different reproductive and feeding stages, as well as that of the pelagic environmental conditions. In particular, the relatively low n-3 PUFA levels (which are crucial for reproductive success) found in spawning sardines compared to spawning anchovies indicate a poorer reproductive health status of sardine. By comparing the current total lipid content values with those recorded in other Mediterranean and North Atlantic areas, and, others from more than ten years ago, in the adjacent area of the Gulf of Lion, our study reveals the persistent poor condition of sardine and anchovy in the northern Catalan Sea. Furthermore, the low levels of diatom fatty acid markers observed throughout the spawning and non-spawning seasons in both sardine and anchovy, indicate a diet poor in diatoms. Moreover, the results indicate that it is very unlikely that parasitism is a significant factor in the decline in condition of sardine and anchovy in the northern Catalan Sea. In fact, the results suggest that the current poor condition of sardine and anchovy in the northern Catalan Sea has been exacerbated by a decrease in plankton productivity and/or a shift in the taxonomic composition of phytoplankton communities, adding to the ongoing effects of overfishing.

## Introduction

Substantial declines in the stock size, mean body size and/or condition of European sardine (*Sardina pilchardus*) and European anchovy (*Engraulis encrasicolus*) have been observed in the north-western Mediterranean Sea since 2009 (Van Beveren et al. 2014; Brosset et al. 2015, 2016a; 2017; Ferrer-Maza et al. 2016, Albó-Puigserver et al. 2017, 2019; Saraux et al. 2019), resulting in profound changes in the structure of the stocks and a major decline in the landings and fishing activity (Coll and Bellido 2019; Brosset et al. 2017; Saraux et al. 2019). Similar negative trends in the body condition of sardine and anchovy have been documented in other northern areas of the Mediterranean Sea (Brosset et al. 2017) and for sardine in the Bay of Biscay in the North Atlantic (Veron et al. 2020). The current status of sardine and anchovy stocks is worrying as these small pelagic species are not only important to fisheries, they are also important from an ecological point of view, as they have a central place in the food web as forage species (Saraux et al. 2019; Albó-Puigserver et al. 2019). Forage fish play a fundamental role in marine trophodynamics because they uptake the energy available from low-level plankton and provide higher-order predators, including marine mammals, seabirds, large piscivorous fish and humans, with a highly nutritious and energetic food source (Cury et al. 2000). Hence, changes in body condition in these small pelagic fish can have important implications for the whole ecosystem structure (Pethybridge et al. 2014; Albó-Puigserver et al. 2017, 2019; Saraux et al. 2019).

Overfishing, climate change, diseases, predation by large fish such as tuna, and competition between pelagic organisms for the zooplankton they feed on, have all been suggested as factors to explain the decline in abundance and mean weight of sardine and anchovy populations in the Gulf of Lion. It seems, however, that the combined effects of poor condition, slower growth and the disappearance of older and larger individuals mediated by potential changes in food availability have been the major causes (Saraux et al. 2014, Van Beveren et al. 2014; Saraux et al. 2019). In the NW Mediterranean, anchovy feed on zooplankton (particularly large copepods) whereas sardine feed on both zooplankton (mainly large copepods) and phytoplankton (mainly diatoms) (Plounevez and Champalbert (2000), Costalgo and Palomera (2014); Le Bourg et al. (2015). However, recent studies have suggested a shift in the diet of sardines in the Gulf of Lion from larger mesozooplankton (with a high proportion of cladocerans) before 2008 to smaller prey (copepods, suspected to be less nutritious) in the early 2010s (Zarubin et al. 2014; Brosset et al. 2016b). Furthermore, an experimental study carried out in the Gulf of Lion showed that food size is as important as food quantity for body condition, growth and total lipids of sardines (Queirós et al. 2019). A combination of pollution and sea warming may have resulted in a long-lasting domination of smaller, lower-energy plankton in this region, which could be extremely detrimental to sardine populations (Queirós et al. 2019). Overall, plankton composition, concentration and size seem to play a key role in determining the condition of small pelagic fish as other studies have shown: anchovy in the Strait of Sicily (Basilone et al. 2004, 2006) and in the Adriatic (Zorica et al. 2013), sprat in the Black Sea (Shulman et al. 2005) and sardine in the Bay of Biscay (Veron et al. 2020). In the Bay of Biscay, the decline in body condition in sardine since the late 2000s had no apparent link with fishing pressure but instead was linked to trophic responses involving a potential shift in the timing of the secondary production and/or the quality of the food (Veron et al. 2020).

Assessing fatty acid composition in forage fish is seen as an ideal way to understand variability in their population dynamics (Shulman et al. 2005; Litzow et al. 2006; Lloret et al. 2014; Pethybridge et al. 2014; Keinänen et al. 2017). In addition, fatty acid composition can be used to monitor energy availability and energy transfer in a food web, because it is known to reflect the fatty acid content of the fish diet, and, ultimately, of local phytoplankton (St. John and Lund 1996; Litzow et al. 2006), and to determine the flow-on effects of these observed changes to their predators, because lipid content in forage fish is likely to have a large influence on higher-order secondary production (Lloret et al. 2014; Pethybridge et al. 2014; Keinänen et al. 2017).

Fatty acids are relevant from a nutritional point of view because they serve as substrates for a number of important metabolic energy and maintenance processes that underlie essential life history traits of fish, such as reproduction, growth and development. Fatty acids are the most important components of lipids, defining their energy value and forming the structural–metabolic “skeleton” of cellular and subcellular membranes. In particular, polyunsaturated fatty acids (PUFAs) – among which are the omega 3 fatty acids, such as eicosapentaenoic acid (EPA) and docosahexaenoic acid (DHA), and the omega 6 fatty acids, such as arachidonic acid (ARA) – are fundamental components of membranes and are regarded as essential for ion transport and for regulating the viscosity of membranes, as they provide osmotic and electrolytic homeostasis and membrane permeability (Lloret et al. 2014). In addition, PUFAs have been identified as a major dietary factor in determining successful reproduction of fish, being crucial for the future requirements of the progeny (Tocher 2003; Lloret et al. 2014). They affect hatching success and viability of larvae because they are especially important in the development of larval activity and vision, as they accumulate in muscle, retinal rhodopsin, and brain tissue of larvae and provide them with a better orientation during feeding (Tocher 2003; Lloret et al. 2014). EPA and ARA are precursors of prostaglandins, which have a role in final oocyte maturation and ovulation (Lloret et al. 2014). Selective retention of DHA and ARA in ovaries during ovarian maturation occurs in species such as cod (Røjbek et al. 2012). In the case of sardine and anchovy, there is evidence of the importance of fatty acids in their reproduction success. For example, a significant variation in the EPA and ARA concentration of Iberian sardine oocytes was found to be caused by parental effects, with the amount, and particularly the composition, of the fat reserves that sardines are able to accumulate prior to the spawning season having a marked effect on the quality of the eggs produced during the spawning season (Garrido et al. 2007). Hence, from an eco-physiological perspective, assessing PUFAs is one of the best ways to test the effects of lipid reserves on the reproductive success of small pelagic fish (Lloret et al. 2014). However, the majority of marine fishes do not possess the ability to synthesize PUFAs themselves: in pelagic ecosystems, they are mostly produced only by phytoplankton and are transferred up the food webs; hence, they are considered to be essential fatty acids (EFAs; Dalsgaard et al. 2003; Lloret et al. 2014).

Furthermore, determining fatty acid profiles can help in monitoring ecosystem dynamics in the face of global climate change, reflecting baseline food web dependencies (Auel et al. 2002; Dalsgaard et al. 2003). In a changing ocean, studies of the fatty acid profiles of forage fish, complemented with other physiological measures such as oxidative stress balance, could help reveal shifts in primary productivity and consequently lead to a system-level understanding of marine trophodynamics (Litzow et al. 2006; Pethybridge et al. 2014; Queirós et al. 2019).

Along with food availability and reproduction, parasitism has also been identified as a factor affecting the body condition of several fish species in the Mediterranean (e.g. Lloret et al. 2012; Ferrer-Maza et al. 2014, 2015; 2016; Serrat et al. 2019). However, to our knowledge, only two studies have looked into the effects of parasites on the lipid content of small pelagic fish in the Mediterranean. The first, Shchepkina (1985), analyzed the lipid concentration in the liver and the white and red muscles of anchovy in the Black Sea and found that specimens that were heavily infected by nematodes showed lower lipid concentrations (especially triglycerides) in their tissues than lightly infected specimens. The second study, by Ferrer-Maza et al. (2016), revealed that certain parasites could be having a negative effect on the energy reserves of anchovy and hypothesized that the differences observed in energy reserves in anchovy could be due to the effect of parasitism rather than reproduction. However, a study from Van Beveren et al. (2016) did not provide evidence of strong pathogenicity from parasites in sardine and anchovy in the Gulf of Lion.

In this context, this study analyses, from an ecological standpoint, the fatty acid composition of sardine and anchovy from the northern Catalan Coast (NW Mediterranean) in different reproductive and feeding stages, in order to assess the links between reproduction, feeding and lipid dynamics in both species. We also evaluate a number of fatty acid trophic markers that have been proposed as candidates for assessing changes in the condition of small pelagic fish related to changes in planktonic productivity. In addition, we compare the results provided in this paper in the northern Catalan Coast with results from other areas to shed light on the current situation in the northern Catalan Sea. Finally, the lipid dynamics are complemented with the analysis of an extensive parasitism data set in order to establish whether or not parasites are in some way responsible for the poor status of these small pelagic species in the study area.

## Materials and methods

### Sampling of individuals

Samples of adult sardines and anchovies caught by purse seines were taken at the ports of Blanes, Roses, Sant Feliu de Guíxols, L’Escala and Palamós in the northern Catalan Sea (Figure 1), during two different periods, corresponding to the two spawning seasons reported for each species: first, the spawning season of sardine in autumn/winter period from November 2018 to January 2019 (Palomera and Olivar 1996; Ganias et al. 2007; Hani et al. 2016); and second, the spawning season of anchovy in the spring/summer period from April 2019 to June 2019 (Palomera 1992. In order to verify reproductive status, half the specimens sampled were assessed for maturity stage via visual inspection of the gonads after dissection. Based on collected commercial data and maturity stage descriptions for anchovy and sardine (ICES 2008), we concluded that our spawning individuals were in the categories *spawning capable* or *spawning* (i.e. in the early and active reproductive periods) whereas non-spawning individuals were in the categories *post-spawning, resting* or *developing*. Henceforth we shall refer to these as “spawning anchovy/sardine” and “non-spawning anchovy/sardine”. Inspecting the gonads of the specimens allowed us to relate lipid dynamics to reproductive cycle more precisely, compared to inferring the reproductive stage of the individuals from the month of capture, as was the case in previous works (e.g. Pethybridge et al. 2014).

**Figure 1.**
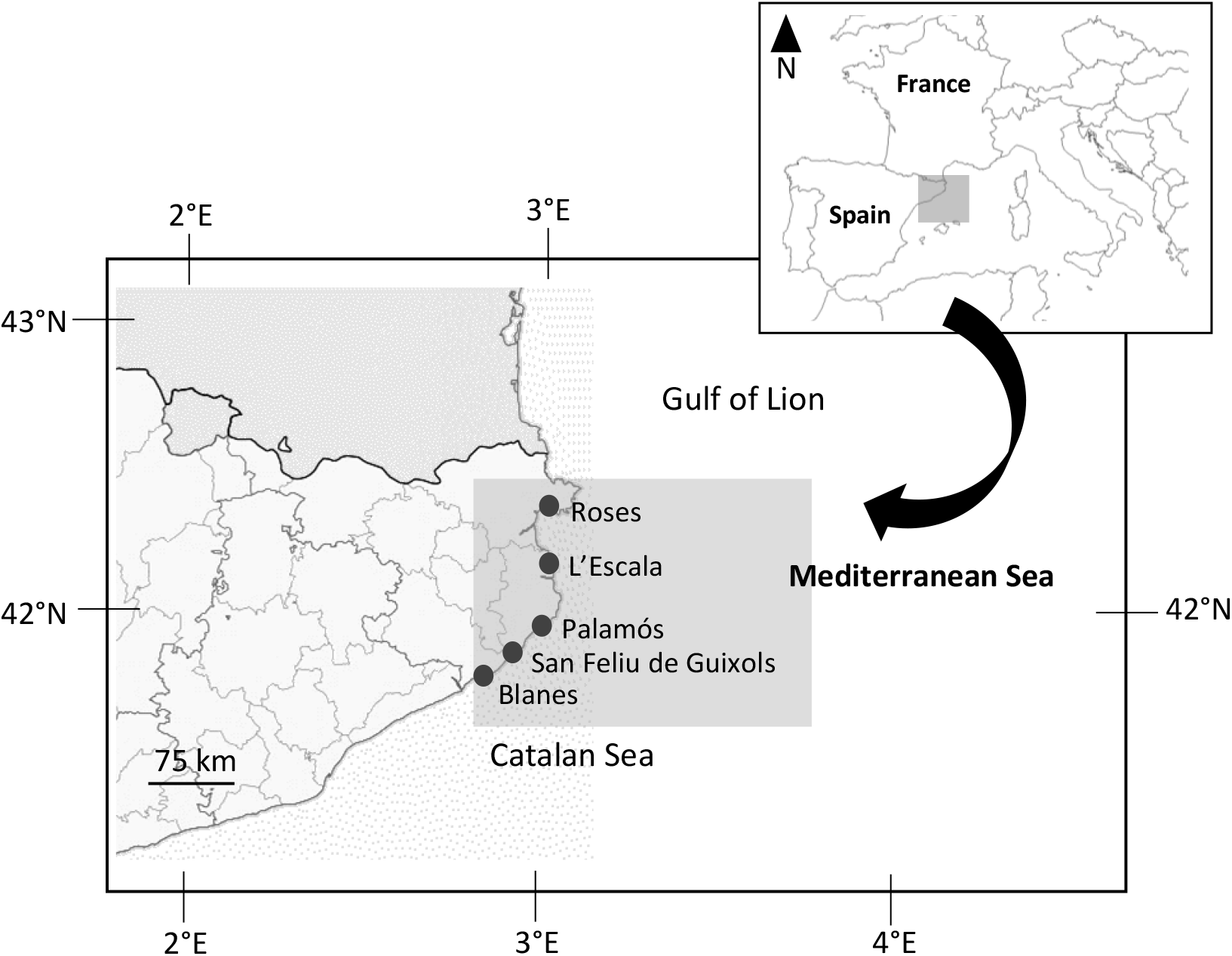
Map of the fishing ports in the northern Catalan Sea (NW Mediterranean) where samples of sardines and anchovies were taken

Sampling was performed on several days each month. Fish catches were grouped in different sample units, which were classified as follows: 10 samples of spawning sardines, 15 samples of non-spawning sardines; 9 samples of spawning anchovies and 13 samples of non-spawning anchovies. Sample units consisted of between 15 and 90 anchovies (similar lengths, randomly selected from the catch) and between 15 and 40 sardines (similar lengths, randomly selected from the catch). Because the effect of sex and length on lipid content and fatty acid levels of sardine and anchovy was not significant (as reported in the Gulf of Lion by Pethybridge et al. 2014), we grouped the specimens together. Individuals were headed and gutted within 24 h of being caught, and the muscle samples were homogenized with a grinder and kept frozen at −80 C until analysis.

### Analysis of total lipid (fat) and fatty acids

The fat, or lipid content and fatty acid composition were determined for the muscle of sardine and anchovy, where both species, and indeed most pelagic fishes, store most of the energy reserves (review by Lloret et al. 2014). The total lipid content (% wet weight) was determined with an automatic Soxhlet extractor (Gerhardt SOX-416 Macro) following ISO 1443:1973 for fat extraction. Ground samples were first hydrolysed with hydrochloric acid (100 ml water+50 ml hydrochloric acid for every 10 grams of sample) and the lipid fraction was extracted by repeated extraction (percolation) with a volume of 150 ml of petroleum ether per 10 grams of sample. This solvent flowed for several cycles through the sample into a glass vitrified capsule (thimble) by distillation. The lipid content in the samples was then calculated by differences in weight.

Fatty acid methyl esters (FAMEs) were analyzed by Gas Chromatography coupled with a Flame Ionization Detector (GC-FID) following ISO 12966-4:2015. First, 30 g of ground sample were extracted with 50 ml of petroleum ether. The extract was then evaporated by means of a Buchi rotary evaporator R-210. FAMEs were prepared by transesterification of the lipid extract, according to ISO 12966-2. FAMEs were analyzed using an Agilent 7693A gas chromatograph coupled to a FID (Agilent Technologies, US). The injection volume of samples and standards was 1μL and the column used was a high-polarity capillary column, BPX 70 (70% cyanopropyl / polysilphenylene-siloxane column, 30 m x 0.25 mm; 0.25 μm film thickness). Initial temperature was 90°C for 1 min, followed by a ramp of 4°C/minute up to 206°C and then another ramp of 20 °C/minute up to 246 °C at which point the temperature was held for 5 min. Detector and injector temperatures were set at 280°C and 260°C, respectively. The whole process lasted 37 minutes, with an air flow of 400 mL/minute, an H2 flow of 30 mL/minute and a Helium flow of 25 mL/minute. Chromatographic peaks were integrated and identified using standard samples (Supelco 37 Component FAME Mix, from Sigma Aldrich). The content of each fatty acid in lipids was expressed as a percentage of the total content of all fatty acids. A total of 24 fatty acids were identified in the total lipid fraction from both species. However, some were detected at such low levels that a cut off point for quantification was set at 0.1% for both fish species. This resulted in the quantification of 16 fatty acids.

### Indices of trophic relationships

In order to assess trophic relationships, we computed the following ratios (Auel et al. 2002; Dalsgaard et al. 2003): palmitoleic acid/palmitic acid (16:1 n-7/16:0; or PO/P) and eicosapentaenoic acid/docosahexaenoic acid (20:5 n-3/22:6 n-3; or EPA/DHA). High values of these ratios indicate a diatom-based diet, whereas low values indicate a dinoflagellate-based diet. This is because among the specific lipid components suggested as suitable for use as trophic biomarkers in the pelagic marine environment (Dalsgaard et al. 2003), diatoms contain high levels of PO, 16:1 n-7 and EPA, 20:5 n-3, whereas dinoflagellates usually contain elevated concentrations of stearidonic acid (18:4 n-3 or SDA) and DHA. Moreover, high EPA/DHA ratios indicate a diet that is predominantly carnivorous (zooplanktivorous), whereas low EPA/DHA ratios indicate a more herbivorous (phytoplanktivorous) diet (Dalsgaard et al. 2003). High EPA/DHA ratios may also be indicative of an important influence of the primary production of cold-diatoms, since cold-water diatoms accumulate especially high amounts of EPA (Falk-Petersen et al. 1998; Scott et al.1999).

### Evaluation of parasitism

To evaluate parasitism, we used the data provided by the Catalan Health Agency gathered from a 6-year program (2002-2007) that monitored parasites in exploited fish species landed in Catalan ports (Servei de Veterinària de Salut Pública, 2007). Samples were collected randomly on a monthly basis by the Agency’s veterinary inspectors at seven of the main fishing ports on the northern Catalan coast. The specimens were caught in the same areas (although in different years) where the specimens used to evaluate fatty acids were caught. In total, 1,269 sardines (measuring between 11 and 31 cm) and 773 anchovies (measuring between 7 and 23 cm) were analyzed for the presence of macroparasites. Immediately after landing, the inspectors recorded the total body length of each specimen and examined them for macroparasites in the gills, skin, fins and intestines using a binocular microscope in facilities at each port. When found, parasites were preserved in a lactophenol solution composed of 1:2:1 lactic acid, glycerol and water. The preserved parasites were then sent to the laboratories of the Catalan Centre of Microbiology where they were identified to the lowest possible taxa possible. 23% of sardine parasites and 16% of the anchovy parasites could be not identified. The prevalence of parasites was calculated as the proportion of fish infected with parasites, whereas the mean intensity of parasitism was calculated as the average number of parasites found in the infected hosts.

### Statistical tests

For each fish species, a one-way ANOVA, considering the spawning and non-spawning period as a factor, was used to examine the existence of significant differences in fatty acid composition and fat content. A post-hoc Tukey HSD was used to identify statistical differences between means. In addition, a Principal Component Analysis (PCA) was carried out to examine the relationships between fatty acid profiles, fatty acid ratios (PO/P and EPA/DHA) and fat content. Furthermore, for each species, the difference in the prevalence of parasites between spawning and non-spawning individuals was tested using a Chi-square 2×2 contingency table. In all cases, the statistical significance was predetermined at P < 0.05. All analyses were performed using JMP13 software (SAS Institute, Cary, North Carolina, USA).

## Results

### Total lipid content (fat) and fatty acid profiles

The values for total lipid content (% wet weight) in the muscle tissue of sardine and anchovy are shown in Table 1. Lipid content was significantly lower in muscle from spawning sardine (mean value, 1.78%) compared to non-spawning sardine (mean, 5.86%). In contrast, lipid content was significantly higher in muscle from spawning anchovy (mean, 2.46%) compared to non-spawning anchovy (mean, 0.89%).

**Table 1.** Total fat (% wet weight) and fatty acid profiles (% of total fatty acids) in the muscle of spawning and non-spawning sardine and anchovy (1)

|  | Sardine |  |  | Anchovy |  |  |
| --- | --- | --- | --- | --- | --- | --- |
|  | Non-spawning | Spawning | Sig. | Non-spawning | Spawning | Sig. |
| Total fat | 5.86 ± 2.14 | 1.78 ± 1.05 | *** | 0.89 ± 0.63 | 2.46 ± 1.49 | ** |
| C14:0 | 7.41 ± 0.90 | 8.07 ± 1.00 | ns | 6.90 ± 2.25 | 6.23 ± 1.74 | ns |
| C16:0 (P) | 23.75 ± 1.63 | 29.49 ± 7.35 | ** | 28.89 ± 6.41 | 21.39 ± 3.50 | ** |
| C17:0 | 0.91 ± 0.16 | 1.44 ± 0.34 | *** | 1.37 ± 0.32 | 0.66 ± 0.44 | *** |
| C18:0 | 4.45 ± 0.63 | 6.59 ± 1.55 | *** | 5.68 ± 1.13 | 2.93 ± 1.42 | *** |
| C20:0 | 0.32 ± 0.12 | 0.43 ± 0.27 | ns | 0.24 ± 0.31 | 2.14 ± 1.22 | *** |
| C22:0 | 0.15 ± 0.09 | 0.16 ± 0.10 | ns | 0.08 ± 0.10 | 0.04 ± 0.09 | ns |
| C24:0 | 0.45 ± 0.45 | 0.15 ± 0.10 | * | 0.34 ± 0.68 | 0.12 ± 0.33 | ns |
| Total SFA | 37.49 ± 2.03 | 46.33 ± 8.76 | *** | 43.50 ± 8.50 | 33.52 ± 5.68 | ** |
| C16:1 n-7 (PO) | 5.59 ± 1.10 | 4.99 ± 1.24 | ns | 4.30 ± 1.33 | 3.84 ± 2.11 | ns |
| C18:1 n-9 | 14.11 ± 1.37 | 12.66 ± 1.65 | * | 11.92 ± 1.73 | 9.69 ± 2.06 | * |
| C20:1 n-9 | 3.57 ± 1.19 | 2.45 ± 1.97 | ns | 0.82 ± 0.95 | 1.52 ± 1.05 | ns |
| C22:1 n-9 | 0.22 ± 0.21 | 0.23 ± 0.21 | ns | 0.10 ± 0.17 | 0.53 ± 0.22 | *** |
| Total MUFA | 23.61 ± 2.23 | 20.33 ± 4.48 | * | 17.15 ± 2.87 | 15.59 ± 4.63 | ns |
| C18:3 n-3 | 1.95 ± 0.64 | 1.40 ± 0.16 | * | 1.07 ± 0.80 | 1.37 ± 0.86 | ns |
| C20:5 n-3 (EPA) | 10.91 ± 0.99 | 7.88 ± 1.42 | *** | 9.02 ± 1.58 | 14.13 ± 2.61 | *** |
| C22:6 n-3 (DHA) | 22.88 ± 2.38 | 20.86 ± 4.17 | ns | 25.24 ± 6.75 | 32.02 ± 7.46 | * |
| Total n-3 PUFA | 35.75 ± 2.70 | 30.13 ± 5.42 | ** | 35.31 ± 8.01 | 47.53 ± 8.64 | ** |
| C18:2 n-6 | 2.26 ± 0.26 | 1.96 ± 0.12 | ** | 1.92 ± 0.50 | 2.43 ± 0.33 | * |
| C20:4 n-6 | 0.89 ± 0.21 | 1.22 ± 0.77 | ns | 2.12 ± 1.21 | 0.97 ± 0.15 | * |
| Total n-6 PUFA | 3.13 ± 0.18 | 3.19 ± 0.75 | ns | 4.03 ± 1.36 | 3.37 ± 0.46 | ns |
| Total PUFA | 38.88 ± 2.75 | 33.32 ± 5.82 | ** | 39.34 ± 8.59 | 50.90 ± 8.89 | ** |
| PO/P | 0.24 ± 0.05 | 0.18 ± 0.07 | * | 0.15 ± 0.05 | 0.18 ± 0.09 | ns |
| EPA/DHA | 0.48 ± 0.07 | 0.38 ± 0.04 | *** | 0.38 ± 0.09 | 0.45 ± 0.11 | ns |

(1) Pairs of means corresponding to spawning and non-spawning fish were compared and those that were significantly different are identified with the symbols \*\*\*, \*\* and \*, showing significance levels of $P < 0.001$ , $P < 0.01$ and $P < 0.05$ , respectively. ns means not significant. Values in this table correspond to means ± standard deviation.

The fatty acid compositions of the total lipid fraction (from the muscle in all cases) of both sardines and anchovies are presented in Table 1.

Saturated fatty acids (SFAs): between 37.49% and 46.33% of the total fatty acids in sardine and between 33.52% and 43.50% in anchovy were SFAs. The most abundant SFA in sardine and anchovy (spawning and non-spawning) was C16:0. Significant differences in the proportion of certain fatty acids between spawning and non-spawning fish were observed. The proportion of C16:0, C17:0, C18:0 – as well as total SFAs – was significantly higher in spawning sardine than in non-spawning sardine, while the reverse was true for C24:0, which was significantly higher in non-spawning sardine. In the case of anchovy, the proportion of C16:0, C17:0, C18:0 – as well as total SFAs – was significantly lower in spawning anchovy than in non-spawning anchovy, while the reverse was true for, in this case, C20:0, which was significantly higher in non-spawning anchovy (Table 1).

Monounsaturated fatty acids (MUFAs): between 20.33% and 23.61% of total fatty acids in sardine muscle and between 15.59% and 17.15% of total fatty acids in anchovy muscle were MUFAs. The most abundant MUFA in sardine and anchovy (spawning and non-spawning) was C18:1n-9. The proportion of C18:1n-9 and total MUFA was significantly lower in spawning sardines than in non-spawning sardines. In the case of anchovy, the proportion of C18:1n-9 was also significantly lower in spawning anchovy than in non-spawning anchovy, but the reverse was true for the proportion of C22:1n-9, which was significantly higher in non-spawning anchovy (Table 1).

Polyunsaturated fatty acids (PUFAs): between 33.32% and 38.88% of the total fatty acids in sardine muscle and between 39.34% and 50.90% in anchovy muscle were PUFAs, most of which were n-3 PUFAs (which comprised between 30.13% and 35.75% of total fatty acids in sardine, and between 35.31% and 47.53% in anchovy). The main differences between the proportion of PUFAs in spawning and non-spawning individuals of both species involve n-3 PUFAs. Among the PUFAs, C22:6 n-3 (DHA) was present in the highest proportion in both species and in both spawning and non-spawning individuals. Significantly lower proportions of total PUFA, C20:5 n-3 (EPA), C18:3 n-3, C18:2 n-6 and n-3 PUFA were found in spawning sardines compared to non-spawning sardines; whereas, significantly higher proportions of C18:2 n-6, EPA, DHA and n-3 PUFA were found in spawning anchovy than in non-spawning anchovy. Only the proportion of C20:4 n-6 was found to be significantly lower in spawning anchovy than in non-spawning anchovy (Table 1).

### Principal component analysis (PCA)

A PCA was performed to examine the variation in fatty acid composition between the two fish species and period of spawning, and to identify the fatty acids most responsible for this variation. The first two components of the PCA explained 62.8% of the variance. As shown in Figure 2, Component 1 influences the majority of SFAs and n-3 PUFAs. Component 1 positively influences C16:0, C17:0, C18:0 and total SFA, and the ratios between C16:1/C16:0 (PO/P) and EPA/DHA, which are localized together and in opposite coordinates to C24:0, C20:5 n3, C18:2 n-6 and C22:1 n-9. In addition, C22:6 n-3, total n-3 PUFAs and C20:0 are grouped together and negatively influenced by Component 1. Meanwhile, Component 2 positively influences fat content and most of the MUFAs. Linolenic acid (C18:3 n-3) and C22:0 are also positively influenced by Component 2. Conversely, n-6 PUFA and C20:4 n-6 are negatively influenced by Component 2.

**Figure 2.**
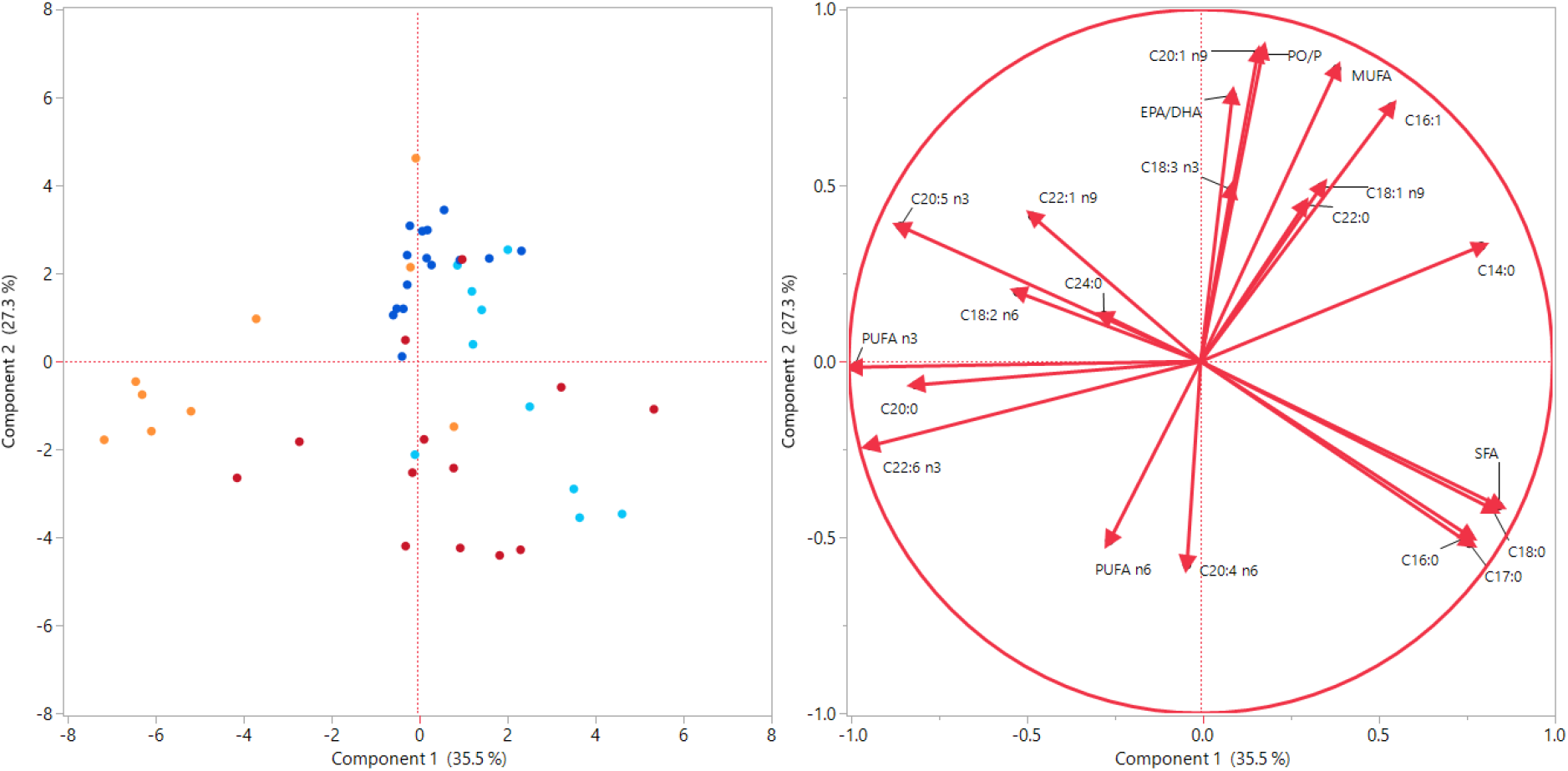
Plots of scores (non-spawning sardine = dark blue; spawning sardine = light blue; non-spawning anchovy = red; spawning anchovy = orange) and loadings of the Principal Component Analysis of fish species fatty acid composition.

In general, the two PCA components allow the variability of fish species to be explained by the period of spawning. Component 1 is associated with the PO/P and EPA/DHA ratios and mainly explains the variability in the fatty acid profile of anchovy due to the spawning period. Accordingly, it seems possible to separate spawning anchovies from non-spawning anchovies by the increase in EPA and DHA (as well as total n-3 PUFAs) and C20:0 and the decrease in the proportion of SFAs with a chain length of up to 18 carbons. Non-spawning sardines are also separated from non-spawning anchovies mainly due to the n-6 and n-9 fatty acid series (Component 2). Similarly, SFAs and n-3 PUFAs help to differentiate between spawning sardines and spawning anchovies (Component 1).

### Parasitism

All the parasites identified in sardines and anchovies were nematode larvae. The results of the Chi-square tests for each fish species showed that the differences in the prevalence of parasites between spawning and non-spawning sardines and anchovies were insignificant. Therefore, the prevalence by species is presented for all individuals (spawning and non-spawning) taken together. Of all the dissected sardine specimens, 7.88% were infected with at least one nematode, with an intensity that ranged between one and three parasites (mean intensity=1.15). *Hysterothylacium* sp was the most frequent parasite, comprising 75.00% of the total nematodes identified, followed by *Anisakis* sp (22.92% of the total). Of all the dissected anchovy specimens, 12.16% were infected with at least one nematode, with an intensity that ranged between one and four parasites (mean intensity=1.10). Again, *Hysterothylacium* sp was the most frequent parasite, comprising 70.40% of the total nematodes identified, followed by *Anisakis* sp (28.61% of the total).

## Discussion

Our results provide new insights into lipid changes in sardine and anchovy that will contribute to our understanding of the physiology and ecology of these small pelagic species in the Mediterranean Sea in the face of changing environmental conditions.

### Seasonal variation in total lipid content and fatty acid profile in relation to reproduction and feeding cycles

First, this study demonstrates seasonal variations in total lipid content between spawning and non-spawning sardine and anchovy in the northern Catalan Sea, and these variations are linked to the different reproduction and feeding strategies of the two species. For sardine, our study found the lowest total lipid content values during the spawning season, i.e., in the autumn-winter period of low food (plankton) availability; for anchovy, the highest values were found during the spawning season, i.e., in the spring-summer period of high food (plankton) availability. Similar patterns of seasonal variability in total lipid content have already been reported in other studies (Ganias et al. 2007; Sánchez et al. 2013; Pethybridge et al. 2014; Ferrer-Maza et al. 2016; Albo Puigserver, 2019) and are in consonance with the breeding strategy of each species: sardine has been described mainly as a capital breeder, relying on energy stores accumulated prior to reproduction, whereas anchovy has been described mainly as an income breeder, relying on an abundant food source during their spawning phase (García and Palomera 1996, Somarakis et al. 2004; McBride et al. 2015; Brosset et al. 2016b).

Second, our study has been able to link the fatty acid composition of sardine and anchovy in the northern Catalan Sea with the reproductive and feeding cycle of these species. In the case of sardine, and in line with the total lipid content data, levels of total MUFAs and total PUFAs were highest during the non-spawning season during which feeding intensity is high, in consonance with capital breeding strategy. In the case of anchovy, and also in line with the total lipid content data, the total PUFAs were significantly higher during the spawning season, during which the phytoplankton are in maximum supply, in consonance with its income breeding strategy. However, while the differences in MUFAs between spawning and non-spawning anchovies were not significant, the total SFA values in both species displayed the opposite pattern to that of total lipid with significantly higher values in spawning sardine and in non-spawning anchovy.

The reproductive and feeding cycles of these two fish species explain the variability in the fatty acid composition that can be differentiated by means of a PCA. Similar findings have been reported for the fatty acid composition of sprat, sardine and anchovy collected in Gulf of Lion (Pethybridge et al. 2014). These authors reported that C14:0, C16:0, C16:1 n-7, C18:1 n-9, EPA and DHA are crucial for explaining the variability in the fatty acid composition of these fish species which is very consistent with our findings (Figure 2). It is also worth noting that the PO/P and EPA/DHA ratios together with the proportions of C18:1 n-9, and, to a lesser extent, EPA and C14:0, can help to explain feeding habits. It appears, therefore, that the sardine’s diet during spawning is more carnivorous (zooplanktivorous) and less herbivorous (phytoplanktivorous) than it is during non-spawning. However, these indices lack significance in the case of anchovy which suggests that it has different feeding habits compared to sardine, supporting the hypothesis that sardine and anchovy probably do not compete strongly for food resources (Chouvelon et al. 2015). As shown in the PCA, higher percentages in DHA seem to discriminate anchovies during spawning when their diet may be richer in dinoflagellates and, in general, phytoplankton.

Our findings also suggest that, in general, a higher proportion of MUFAs is associated with fattier fish, which may explain the relatively lower proportion of very long chain n-6 PUFAs, with the exception of the precursor of n-6 fatty acid series, linoleic acid (18:2 n-6). On the other hand, the increase in very long-chain n-3 PUFAs seems to be at the expense of SFAs containing up to 18 carbon atoms. SFAs and MUFAs are major sources of metabolic energy in fish (particularly, C16:0, which is a predominant source of potential metabolic energy during growth and ovary development; Henderson et al. 1984), and can be synthesized by the fish themselves. There are, therefore, complex trade-offs between reproduction, growth and basal energy that cannot be fully explained in our study.

### The relevance of PUFAs

We shall now turn our attention to the PUFA levels, bearing in mind that they may provide the only outputs that can be easily interpreted from a physiological point of view. In particular, n-3 PUFAs have been identified as a major dietary factor determining successful reproduction in fish, as they are crucial for the future requirements of the progeny (Tocher 2003, Lloret et al. 2014). There is a high requirement for n-3 PUFAs in the developing eggs and larvae of fish because of their preponderance in neural and visual tissues, which predominate in the early stages of development (Bruce et al. 1999). Hence, any deficiency in these particular fatty acids can cause abnormalities in the neural system and may affect the success of larvae as visual predators at the onset of first feeding (Bell and Sargent 1996). In fact, anchovy larvae in the NW Mediterranean contain a high proportion of PUFAs (Rossi et al. 2006).

The variation in total PUFA levels found in sardine and anchovy in the northern Catalan Sea was mostly due to variations in the levels of highly unsaturated fatty acids, namely the n-3 fatty acids, EPA and DHA. Although the relative proportion of n-3 PUFAs in the fatty acids of non-spawning sardine and non-spawning anchovy was quite similar (about 35%), the relative proportion in spawning sardine was much lower (about 30%) than in spawning anchovy (47%). If we consider the significant relationships found between n-3 PUFAs in female muscle and oocytes of sardine in the North Atlantic, and the relationship between female diet – in particular, plankton availability immediately before and during the spawning season – and the quality of offspring produced by sardine (Garrido et al. 2007), then we can surmise that the relatively low proportion of n-3 PUFAs in spawning sardines in the northern Catalan Sea indicates a poorer reproductive status of this species than that of anchovy. It must be also taken into account that sardines have a lower degree of trophic plasticity than anchovies, both in terms of feeding areas and in the size of the zooplanktonic prey consumed (Chouvelon et al. 2015) and that Van Beveren et al. (2016) revealed elevated quantities of macrophage aggregates in sardines in the Gulf of Lion indicating stress on the fish that might potentially be related to starvation. In the following section we address the issue of the challenging food supply over time in more detail

### What do fatty acids tell us regarding the current status of sardine and anchovy stocks under challenging environmental conditions?

In order to understand the challenges facing small pelagic fish in the Mediterranean, and particularly that of sardine, we shall now discuss how the fatty acid profiles, and the ratios computed, help to explain the potential causes behind the current status of the stocks. In our study, low diatom markers were present throughout the spawning and non-spawning seasons of sardine and anchovy. The low (< 0.50) ratios of PO/P, and EPA/DHA for both sardine and anchovy during the non-spawning periods support the hypothesis of a diet for both species that is not predominantly based on diatoms. This is in contrast to the situation ten years ago (2010 and 2011) in the adjacent waters of the Gulf of Lion, where for all seasons, these ratios indicated a predominantly diatom-based diet for both species (Pethybridge et al. 2014). In fact, studies on stomach analyses of sardine in the Gulf of Lion at that time (2011-2012) showed a higher proportion of diatoms in the diet compared to dinoflagellates, a situation that was more accentuated during summer when diatom abundance was usually high after the spring bloom (Le Bourg et al. 2015; Leblanc et al. 2003).

Furthermore, the relatively low PO/P and EPA/DHA ratios of non-spawning sardine in the northern Catalan Sea compared to the ratios observed in other Mediterranean and North Atlantic areas indicate that, in the Catalan Sea, the proportion of diatoms in the diet of sardine is lower than in other areas (Table 2). Furthermore, the comparatively lower EPA/DHA values and high C16:0 values of non-spawning sardines in the northern Catalan Sea suggest that low-energy phytoplankton is proportionally more important than high-energy zooplankton in the sardine’s diet in the study area compared to other areas. This pattern does not occur in anchovy (Table 3), for which the ratios of PO/P, EPA/DHA and the levels C16:0 in non-spawning individuals from the Catalan Sea are similar to other Mediterranean areas, except in the Black Sea, where higher PO/P, EPA/DHA values and lower C16:0 values are found (Table 3). Notwithstanding these results, the comparison of fatty acid profiles between areas must be taken with caution, because values compared are expressed in % of total fatty acid mass, and it would be much better to compare data on absolute fatty acid content (% body mass) (Litzow et al. 2006).

**Table 2.** Total fat (% wet weight) and fatty acid profiles (% of total fatty acids) during non-spawning periods for *Sardina pilchardus* in the Eastern Algarve waters of the Atlantic Ocean, (Bandarra et al. 2017); the Mediterranean waters of the Adriatic Sea (De Leonardis and Macciola 2004), the Gulf of Lion (Pethybridge et al. 2014) and the northern Catalan Sea (our study).

| Marine Areas | Eastern Algarve<br>October 2016 | Adriatic Sea<br>January–March and<br>August–October 2002 | Gulf of Lion<br>July 2010 | **This study**<br>**(N Catalan Sea)**<br>**November-April**<br>**2019-2020** |
| --- | --- | --- | --- | --- |
| Total fat | 14.0 | (-) | 18.21 | **5.86** |
| C16:0 (P) | 19.6 | 22.3 | (-) | **23.75** |
| C16:1 n-7 (PO) | 6.6 | 9.2 | (-) | **5.59** |
| **PO/P** | 0.34 | 0.41 | (-) | **0.24** |
| C20:5 n-3<br>(EPA) | 13.6 | 6.5 | (-) | **10.91** |
| C22:6 n-3<br>(DHA) | 14.8 | 11.3 | (-) | **22.88** |
| **EPA/DHA** | 0.92 | 0.57 | (-) | **0.48** |

(-) means no data available.

**Table 3.** Total fat (% wet weight) and fatty acid profiles (% of total fatty acids) during non-spawning periods for *Engraulis encrasicolus* in the Mediterranean Sea (Eastern Mediterranean, Black Sea, Oksuz et al. 2009; Tyrrhenian Sea, Adriatic Sea and Ionian Sea, Roncarati et al. 2012; Gulf of Lion, Pethybridge et al. 2014 and northern Catalan Sea (our study).

| Marine Areas | Eastern<br>Medit.<br>December<br>2007 | Black Sea<br>January 2009 | Tyrrhenian Sea<br>May 2007-2008<br>November 2007 | Adriatic Sea<br>May 2007-2008<br>November 2007 | Ionian Sea<br>May 2007-2008<br>November 2007 | Gulf of Lion<br>March 2011 | **This study**<br>**(N Catalan Sea)**<br>**December-June**<br>**2019-2020** |
| --- | --- | --- | --- | --- | --- | --- | --- |
| Total fat | (-) | 8.85 | 2.27 | 1.81 | 1.91 | 8.19 | **0.89** |
| C16:0 (P) | 22.13 | 17.35 | 23.66 | 23.27 | 21.94 | (-) | **28.89** |
| C16:1 n-7 (PO) | 2.61 | 7.93 | 2.95 | 3.17 | 3.71 | (-) | **4.30** |
| **PO/P** | 0.12 | 0.46 | 0.12 | 0.14 | 0.17 | (-) | **0.15** |
| C20:5 n-3<br>(EPA) | 5.65 | 10.92 | 6.13 | 6.25 | 6.49 | (-) | **9.02** |
| C22:6 n-3<br>(DHA) | 33.4 | 15.29 | 28.06 | 27.16 | 25.28 | (-) | **25.24** |
| **EPA/DHA** | 0.17 | 0.71 | 0.22 | 0.23 | 0.26 | (-) | **0.36** |

(-) means no data available

Although our results show relatively similar or higher values of essential fatty acids in non-spawning sardine and anchovy compared to other Mediterranean and Atlantic areas (Tables 2 and 3), when we compare the level of total fat content of non-spawning sardine and anchovy in our study area with that of the Gulf of Lion ten years ago (Pethybridge et al. 2014), we can conclude that the total fat content in the muscle of both species has declined (Tables 2 and 3). This may be related to a decrease in primary production, since a recent study in a coastal area of the northern Catalan Sea showed that most of the phytoplankton groups there presented a decreasing linear interannual trend in abundance, which could be associated with a reduction in nutrient availability (Nunes et al. 2018). Furthermore, the total lipid content of non-spawning sardines and anchovies in the northern Catalan Sea is also much lower than in the other areas (Tables 2 and 3) suggesting that sardine may even be forced to rely on direct food intake to acquire enough energy during the spawning period.

Taking all these results together, it appears that a decrease in plankton productivity and/or a shift in the taxonomic composition of phytoplankton communities may have occurred in the northern Catalan Coast in the last decade, although further investigations are needed to confirm this statement as the studies carried out so far remain scarce (Thiébault et al. 2016). The results from our study support the hypothesis proposed by Saraux et al. (2019) that changes in plankton availability/diversity may be a factor in the poor condition observed in small pelagic fish in the NW Mediterranean. By altering the pelagic environment, climate change may be altering the composition and distribution of plankton species, as well as their importance in the food web, with higher temperatures favouring the smallest components of the plankton, thus strengthening microbial loop activity (Thiébault et al. 2016). As increasing temperature favours planktonic organisms of smaller size, climate change may particularly affect the condition of sardine, as recent results revealed that food size (without any modification of its energy content) is as important as food quantity for the body condition, growth and reserve lipids of sardine (Queirós et al. 2019). Furthermore, fluctuations in the diatom/dinoflagellate ratio may have ecosystem-wide consequences for the transfer of energy and matter to higher trophic levels considering the importance of both groups in the trophic chain of many seas (Wasmund et al. 2017). Although EFA availability is generally high in marine ecosystems, it is also highly variable, implying that, occasionally, EFA availability is limited (Litzow et al. 2006). In the Pacific Ocean, for example, it seems that climate-mediated changes in the availability of EFA were behind the changes in lipid content of different fish communities, supporting a growing consensus that EFA availability may influence trophic structure in aquatic ecosystems (Litzow et al. 2006).

### Parasitism

Our results on parasitism showed that the metazoan parasite fauna of sardine and anchovy in the northern Catalan Sea is dominated by nematode larvae. However, considering the low prevalence and low intensity of parasites in sardine and anchovy in the study area, our results are in line with Van Beveren et al. (2016) and Saraux et al. (2019) both of which concluded that it is very unlikely that parasites, or any other pathogenic agent, are root causes of the drastic population modifications observed in sardine and anchovy in the NW Mediterranean. For sardine, the study by Van Beveren et al. (2016) found no link between sardine condition and a wide range of potential pathogens, including parasites (although only microparasites were found), viruses and bacteria; in other words, no strong indications of pathogenicity were found. In contrast, for anchovy, a study by Ferrer-Maza et al. (2016) showed that certain species of parasites had a negative effect on female egg production and lipid content in smaller individuals. However, there is a wide range of research available concerning the nematodes that infect anchovy and sardine in the Mediterranean (e.g. Rello et al. 2009; Gutiérrez-Galindo et al. 2010; Cavallero et al. 2015; Zorica et al. 2016) and, in general, such studies have reported a relatively low prevalence of nematodes in these two species, ranging from 0 to 25%, with low values for overall prevalence when data from different studies were pooled (<3% in the case of Anisakids in the meta analysis conducted by Colombo et al. 2016). In a study by Rello et al. (2008), *H. aduncum* was the only Anisakid parasite found in sardines from the southern and eastern coasts of Spain, with a total prevalence of 11.85%. Although it is true that prevalence may change depending on season, parasite species and area (Mladineo et al. 2012; Zorica et al. 2016), such low values support the idea that parasites are probably not responsible for the poor status of sardine and anchovy stocks in the Mediterranean.

Our results contribute to the literature on parasitic infestation of sardine and anchovy in the northern Catalan Sea because, for the first time, we used extensive data from a monitoring programme covering many ports and years and seasons. However, it must be noted that other metazoan parasites (such as Digenea and Cestoda) are found in particular organs, such as pyloric caeca and the stomach, which are difficult to evaluate in the port and may have gone undetected by the veterinarian inspectors involved in the programme. It must also be noted, however, that inside the musculature of anchovy, the number of parasites is negligible (Ferrer-Maza et al. 2016). Overall, the evaluation carried out by the inspectors in the frame of the monitoring programme in the northern Catalan Sea ports provide good estimates of parasitism in this region, despite the fact that monitoring parasite infestations in fish is particularly challenging due to the complex interactions among hosts, parasites and the environment, and the existence of many zero-values (Helland et al. 2015).

## Conclusion

The results from this study provide evidence that fatty acid levels are good indicators not only of the body condition at different reproductive and feeding stages of small pelagic fish, but also of pelagic environmental conditions in the context of global change. The study contributes to our understanding of the trade-off between condition, reproduction and feeding in sardines and anchovies through the analysis of variations in the profiles of fatty acids that are crucial in key life history traits of these forage species, particularly reproductive success. In our study, low diatom markers were observed throughout the spawning and non-spawning seasons of sardine and anchovy, indicating a potential shift in the diets of sardine and anchovy in the area from diatoms to dinoflagellates. Overall, these results indicate that a decrease in plankton productivity and/or a shift in the taxonomic composition of phytoplankton communities may have occurred in the northern Catalan Sea in the last decade. Feeding conditions in spring and summer appear to be a key factor in determining not only total lipid content but also the levels of specific fatty acids of sardine and anchovy in the NW Mediterranean. According to our results, it would appear that, based on total lipid content and fatty acid distribution, sardine and, to a lesser extent, anchovy, in the northern Catalan Sea are currently in poor condition and malnourished. In order to confirm this trend and to confirm any potential limitations in PUFA/EFA availability, further studies on the fatty acids of small pelagic fish in the area will be needed. Because lipid biochemistry is complex, more research into the nature of the PUFA/EFA requirements is needed before the ecological implications can be elucidated.

Our results also support the idea that it is very unlikely that parasites are a root cause of the decline in the condition of sardine and anchovy in the NW Mediterranean, but any increase in parasitism could instead be the result of qualitative and/or quantitative modifications in planktonic production leading to fish in poorer condition, particularly sardines. Further studies on the planktonic composition and its evolution in the Mediterranean Sea are needed to improve our understanding of the impacts of changing food quantity and quality on the condition of these small pelagic fish, which may have consequences not only for fisheries but also for the whole trophic chain – considering their importance as forage species. Furthermore, bearing in mind that, in the Catalan Sea, which is part of GSA6, the sardine and anchovy stocks are considered to be overexploited according to STECF assessments (STECF 2016), the poor condition status of these species adds to the worries posed by the impact of fishing activity. This is more evident for sardine, which according to assessments made by GFCM (2017) has seen a negative trend in landings and acoustic biomass estimates since 1994 in GSA 6.

## Acknowledgements

This study was supported by FLAG / GALP-Costa Brava projects Ref. ARP014/17/00007 and Ref. ARP014/17/00054 (financed by the European Maritime and Fisheries Fund – EMFF and the DG Fisheries of the Catalan Govt.). The authors thank A. Rossell, C. Canals and two undergraduate students of the University of Girona (Elena and Cristina) for their help processing samples, as well as to all the fishermen and fishmongers who collaborated in the study. We also thank Dr. Marta Estrada for her valuable comments.

